# “Tumor-selective treatment of metastatic pancreatic cancer with an engineered, probiotic living drug”

**DOI:** 10.1101/2024.05.02.592216

**Authors:** Amanda R. Decker-Farrell, Stephen A. Sastra, Tetsuhiro Harimoto, Marie C. Hasselluhn, Carmine F. Palermo, Edward R. Ballister, Michael A. Badgley, Tal Danino, Kenneth P. Olive

**Author notes:** Correspondence (ARDF), (KPO) or (TD).

## Abstract

Pancreatic ductal adenocarcinoma (PDAC) poses significant challenges for effective treatment, with systemic chemotherapy often proving inadequate due to poor drug delivery and the tumor’s immunosuppressive microenvironment. Engineered bacteria present a novel approach to target PDAC, leveraging their ability to colonize tumors and deliver therapeutic payloads. Here, we engineered probiotic *Escherichia coli* Nissle 1917 (EcN) to produce the pore-forming Theta toxin (Nis-Theta) and evaluated its efficacy in a preclinical model of PDAC. Probiotic administration resulted in selective colonization of tumor tissue, leading to improved overall survival compared to standard chemotherapy. Moreover, this strain exhibited cytotoxic effects on both primary and distant tumor lesions while sparing normal tissues. Importantly, treatment also modulated the tumor microenvironment by increasing anti-tumor immune cell populations and reducing immunosuppressive markers. These findings demonstrate the potential of engineered probiotic bacteria as a safe and effective therapeutic approach for PDAC, offering promise for improved patient outcomes.

## INTRODUCTION

Pancreatic ductal adenocarcinoma (PDAC) is the third leading cause of cancer mortality in the US (*1*). Most patients present with advanced or metastatic disease, making systemic chemotherapies a mainstay of treatment (*2*). However, the biophysical properties of the PDAC tumor microenvironment can interfere with effective delivery of both cytotoxic and targeted agents (*3, 4*). Modern immunotherapy also remains ineffective due to immunosuppressive properties of the microenvironment (*5*), highlighting the need for new treatment approaches.

Interestingly, the features that make PDAC difficult to treat with systemic drugs may make them particularly well-suited for a different approach – targeting via engineered bacteria. Bacteria have been previously utilized to provoke anti-tumor immune responses (*6–12*) and have demonstrated clinical success in some cancers (e.g. Bacillus Calmette--Guerin therapy for bladder cancer (*13*)). Since local immunosuppression can limit the effectiveness of immunotherapy in PDAC, we instead propose to engineer bacteria to serve as “living drugs,” leveraging their ability to colonize immune-privileged tumors, replicate, and produce cytotoxic payloads.

Here, we report the use of probiotic bacteria, *E. coli* Nissle 1917 (EcN), engineered to synthesize the pore-forming Perfringolysin O, also known as Theta toxin (Nis-Theta) (*14*) in the Kras^LSL.G12D/+^; P53^LSL.R172H/+^; Pdx1-Cre^tg/+^ (KPC) genetically engineered model of PDAC (*15*). KPC mice are immune-competent and develop pancreatic tumors that feature an expansive stroma, local immunosuppression, and stark chemoresistance. We found that a single dose of Nis-Theta selectively colonized tumor tissues over normal tissues, potentially due to the inaccessibility of immune effector cells into the immunocompromised tumor microenvironment of PDAC. Treatment with Nis-Theta led to a 3-fold increase in overall survival relative to standard chemotherapy. Strikingly, we found that local administration of EcN to primary PDAC lesions was followed by colonization of secondary tumor sites, with no detectable effects on normal tissues. Finally, in addition to the direct anti-tumor cell effect of the toxic payload, our data show that bacterial presence in the tumors coincided with a modest influx of anti-tumor immune cell populations, while simultaneously reducing immunosuppressive markers. Together, these findings demonstrate the potential of engineered probiotic bacteria to safely and selectively colonize PDAC, disseminate to distal malignant sites, and precisely deliver an anti-tumor cytotoxic payload, yielding an efficacious response in a highly chemoresistant preclinical model.

## RESULTS

### Prioritization of toxin payloads

The effective toxicity of anti-cancer peptides can vary significantly across tumor type and delivery method (*16*). To select an effective anti-tumor toxin for PDAC, we first screened nine established anti-cancer peptide payloads (*16*) in 2D cell culture of murine (KPC) and human PDAC cell lines. Lysates from 4 of the 9 toxin-producing strains (Supp Fig 1A) were effective at reducing the viability of nearly all PDAC lines examined: Theta toxin, Magainin, Heat Stable Enterotoxin (HSEnt), and Hemolysin E (HlyE) (Fig 1A). To more effectively model toxin effects on intact tumor tissue, we carried out a secondary screen using PDAC explant cultures derived from pancreatic tumors in KPC mice (*3, 17*). Of the four candidate strains tested, Theta toxin stood out for its ability to induce apoptosis, as measured by cleaved caspase 3 (CC3) staining, compared to a GFP-expressing control strain (Fig 1B, Supp Fig 1B). Therefore, we elected to develop a therapeutic bacterial strain designed to deliver Theta toxin to PDAC tumors *in vivo*.

**Fig. 1.**
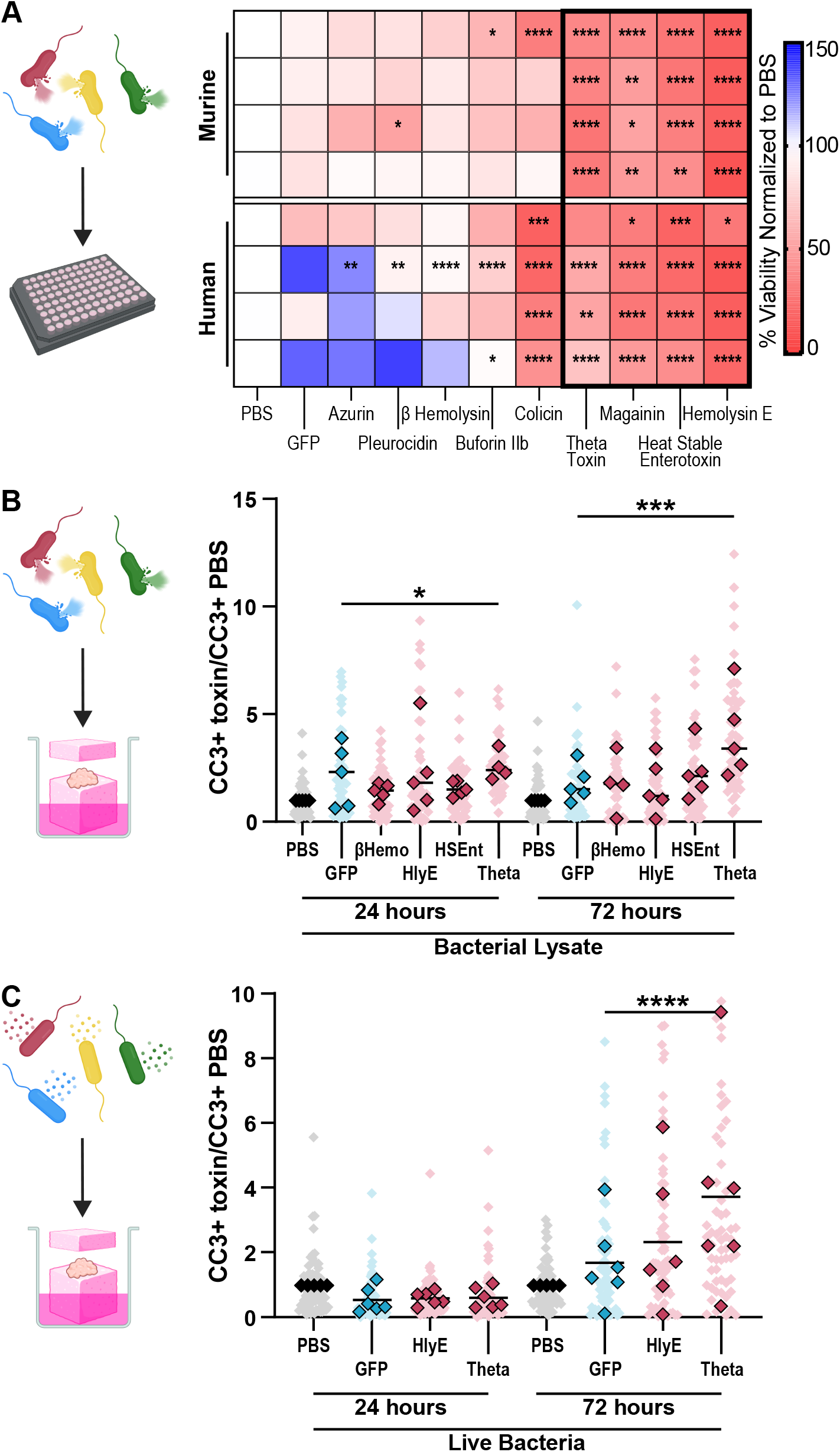
Bacterially expressed Theta toxin induces apoptosis in 2D and 3D models of PDAC. (**A**) Heatmaps quantifying percent viability of human and murine (KPC-derived) PDAC cell lines treated with toxin-containing bacterial lysates for 48 hours in 96-well plates as determined by Alamar Blue and normalized to corresponding PBS treated samples. Two-way ANOVA (Dunnett’s correction), versus GFP treated samples for each cell line. \**P*<0.05, \*\**P*<0.01, \*\*\**P*<0.001, and \*\*\*\**P*<0.0001. (**B**) Quantification of IHC staining for apoptosis marker, CC3, for KPC PDAC explants co-incubated with toxin-containing bacterial lysates for 24 or 72 hours. (**C**) Quantification of IHC staining for apoptosis marker, CC3, for KPC PDAC explants co-incubated with live, toxin-producing *E. coli Nissle 1917* for 24 or 72 hours. Quantification of IHC staining based on 10-12 fields of view at 40x magnification (light shade), averaged per tumor normalized to PBS value for the corresponding time point (solid shade). Data are presented as the mean ± SD. One-way ANOVA (Dunnett’s correction) versus GFP treated samples. \**P*<0.05, \*\*\**P*<0.001, and \*\*\*\**P*<0.0001.

We engineered acylhomoserine lactone (AHL) inducible Theta toxin production into a probiotic strain of *E. coli*, Nissle 1917 (EcN), known to selectively colonize tumors (*18–21*) and is widely utilized in medical products (*22, 23*). As Theta toxin itself can impair bacterial survival and encourage plasmid curing, we stabilized its expression using the toxin-antitoxin plasmid system (AxeTxe or AT) (*24*) (Supp Fig 1C). Additionally, we used a genomically-integrated, constitutively active *lux*CDABE luminescence operon (Nislux (*19*)) to aid in visualization (Nis-Theta). Western blots for Theta toxin confirmed induction of payload production (Supp Fig 1D). Comparable bacteria strains expressing non-toxic GFP (Nis-GFP) and a Hemolysin E (Nis-HlyE) served as negative and positive controls, respectively. Using live toxin-expressing bacteria, a significant increase in apoptosis was observed in only Nis-Theta treated explants, albeit with delayed kinetics compared to treatment with lysates (Fig 1C, Supp Fig 1E). Based on these results, we proceeded to evaluate Nis-Theta for its utility as an anti-tumor living drug against PDAC.

### Anti-tumor effects of Theta toxin-producing bacteria

To evaluate the safety and efficacy of Nis-Theta, we performed an interventional survival study in genetically engineered KPC mice (*15*), a clinically relevant model of PDAC. Tumor-bearing KPC mice were enrolled into the study upon detection (Fig 2A) of a nascent tumor by 3D high resolution ultrasound (*25*). To enhance tumor colonization, tumor-bearing mice were pretreated with an oral antibiotic and then Nislux strains were administered by ultrasound-guided intratumoral injections (Fig 2B, Movie S1) and monitored twice weekly (Fig 2A) for tumor growth (*25*) and bacterial luminescence (*19*).

**Fig. 2.**
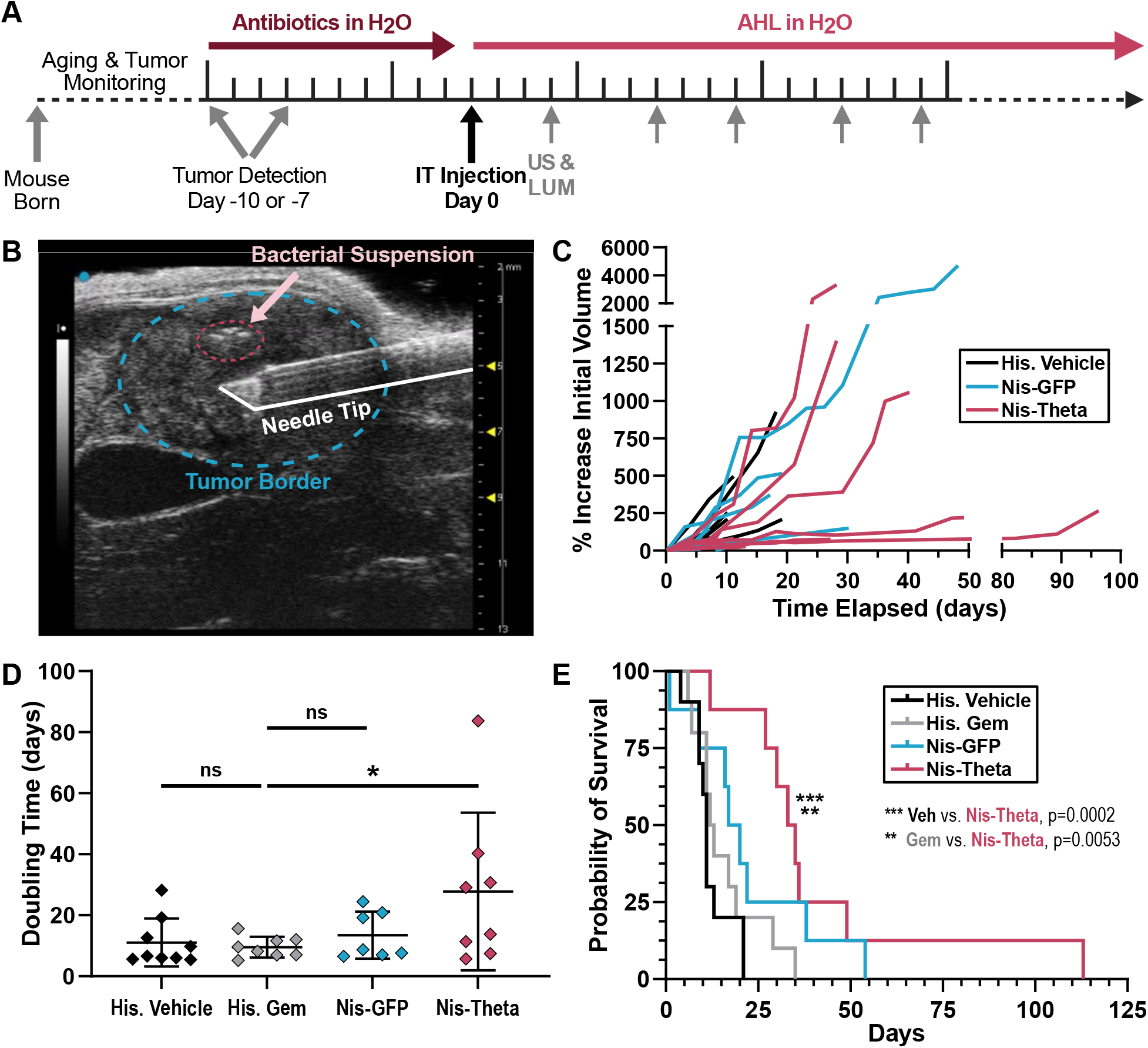
Intratumorally delivered Nis-Theta transiently slows tumor growth and extends survival of KPC mice. (**A**) Diagram of KPC mouse enrollment and treatments with live Nis-GFP or Nis-Theta bacteria (10^7^ bacteria) (n=8 each) for survival study. US = ultrasound, LUM = luminescent imaging, AHL = acyl-homoserine lactone. (**B**) Image of ultrasound-guided intratumoral injection of bacterial suspension into KPC tumor. (**C**) Growth curves of individual tumor volumes as a percentage of the initial tumor volume determined by ultrasound reconstructions. (**D**) Calculated doubling times of reconstructed individual tumor volumes. Data are presented as the mean ± SD. One-way ANOVA (Dunnett’s correction) versus historical vehicle treatment group. (**E**) Kaplan Meier survival curve of KPC mice treated with vehicle (His. Vehicle) (n=10, median 11 days), Gemcitabine (His. Gem) (n=10, median 12.5 days), Nis-GFP (n=8, median 18.5 days), or Nis-Theta (n=8, median 34 days). \*\**P*<0.01, \*\*\**P*<0.001, log-rank.

Tumor growth and overall survival of mice treated with Nis-Theta were compared to contemporaneous and historical controls (*26*). We observed reduced growth rate in Nis-Theta treated tumors (Fig 2C,D) and this was associated with a significant extension of overall survival (Fig 2E).

To further investigate the loss of anti-tumor effects over time, we examined quantity and therapeutic capability of engineered bacteria in primary tumor tissue at time of death. We found no difference in overall colonization among strains but observed a time-dependent decrease in bacterial luminescence (Fig 3A, Supp Fig 2, Supp Fig 3A). Furthermore, all bacteria populations recovered from treated tumors exhibited therapeutic plasmid loss (Supp Fig 3B).

**Fig. 3.**
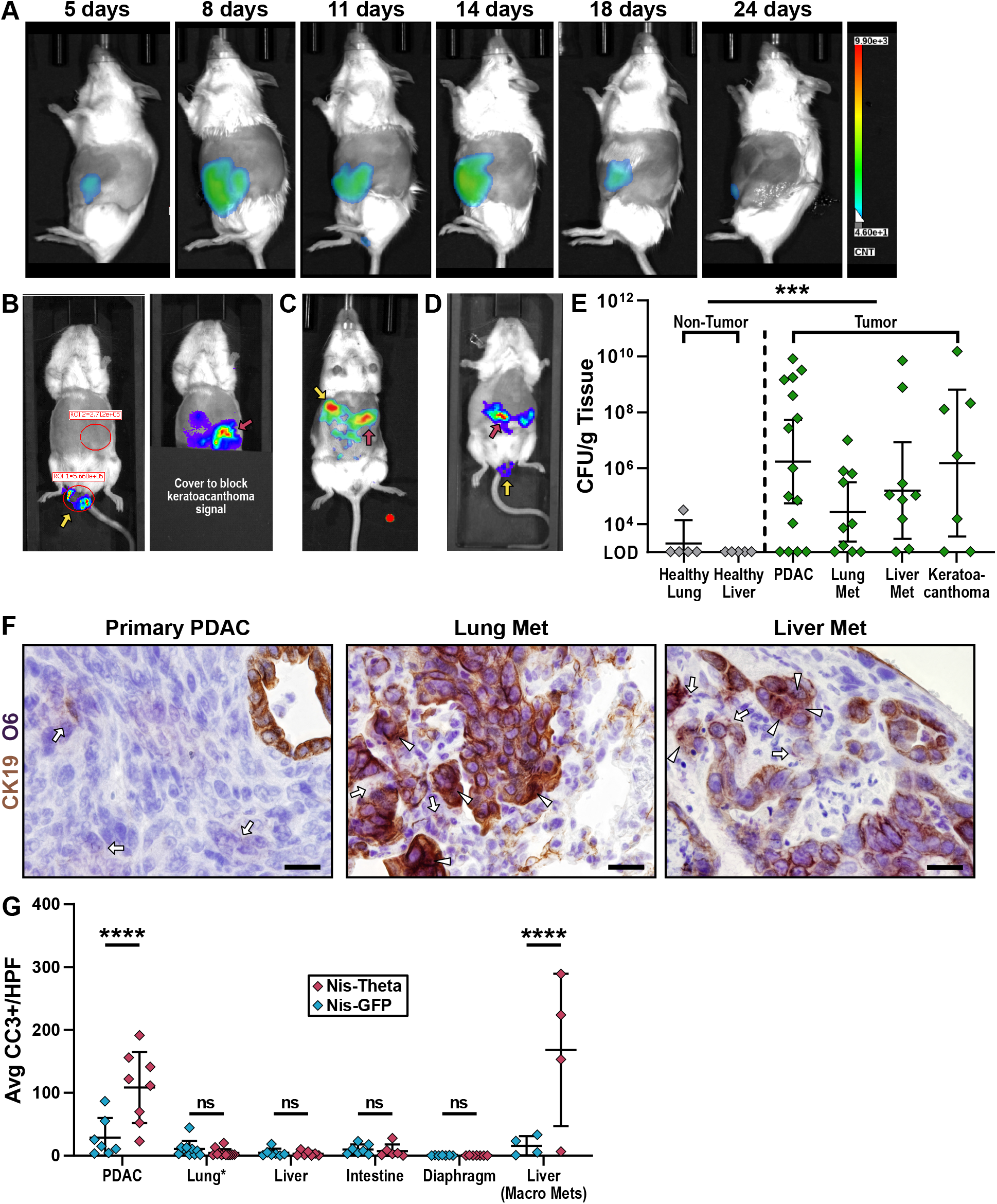
Therapeutic bacteria colonize primary and distant tumor lesions following local, intratumoral delivery. (**A**) Representative time course series of bacterial luminescence in a single mouse through the duration of study treated with luminescent bacteria intratumorally (AMI optical imager, 45 second exposure). (**B**) Bacterial luminescence images of a tumor bearing KPC mouse with an anal keratoacanthoma, following intratumoral injection of the primary PDAC lesion, demonstrating colonization of both the anal keratoacanthoma (yellow arrow) and primary PDAC (red arrow) (IVIS optical imager, 5 minutes exposure). (**C**) Bacterial luminescence image of a tumor bearing KPC mouse following intratumor injection of the primary PDAC lesion, demonstrating colonization of both the primary PDAC (red arrow) and a liver metastatic lesion (confirmed upon necropsy, yellow arrow) (AMI optical imager, 45 second exposure). (**D**) Bacterial luminescence image of a tumor bearing KPC mouse with an anal keratoacanthoma following intratumoral injection of the keratoacanthoma, demonstrating colonization of both the keratoacanthoma (yellow arrow) and primary PDAC (red arrow) (IVIS optical imager, 5 minutes exposure). (**E**) Endpoint biodistribution of Nislux bacteria in peripheral tissues (lung, liver) and tumor tissue (primary PDAC and keratoacanthoma). Peripheral organs were stained with CK19 for metastatic lesions, and divided into healthy (CK19-, gray) or tumor tissue (MET) (CK19+, green). Data are presented as geometric mean ± 95% CI. CFU = Colony Forming Units. \*\*\**P*<0.001, Mann-Whitney, non-parametric t-test. (**F**) Representative images of dual IHC for malignant epithelial lesions (CK19, brown) and Nislux bacteria (O6, purple) demonstrating co-localization of bacteria with metastatic lesions in stromal (arrow) and malignant (arrowhead) compartments. (100x magnification, scale bar = 20 μm). (**G**) Quantification of IHC staining for apoptosis marker, CC3, in peripheral tissues (lung, liver, intestines, and diaphragm) and tumor tissue (primary PDCA and CK19+ liver lesions). Lung* indicates that due to the small size of CK19+ lesions, accurate CC3 data could not be collected solely for metastatic lung tissue, and both CK19+ and CK19-lung tissues are represented in one group. Data were collected from 8-10 fields of view per tissue, of which the averaged values per sample are presented, with bars indicating mean ± SD. Two-way ANOVA (Bonferroni’s correction) of Nis-GFP vs. Nis-Theta for each tissue group. HPF = High Power Field. \*\*\*\**P*<0.0001 and ns, not significant.

### Systemic tumor colonization after local bacterial delivery

Primary pancreatic tumors in KPC mice frequently metastasize to distal sites such as liver metastases, and are often accompanied by synchronous oral or anal keratoacanthomas (*27*). While conducting imaging of KPC animals on the survival study, we observed that intratumoral treatment with Nislux was often followed by colonization of co-occurring distal tumor masses (Fig. 3B,C). To test if bacteria administered to off-site lesions could traffic to primary tumors, we directly injected Nislux into a keratoacanthoma of a KPC mouse with a synchronous PDAC mass. We observed colonization of the PDAC within 4 days (Fig. 3D), indicating that these bacteria can quickly traffic to and colonize distal tumor sites.

As spread to distant locations following local administration suggested systemic bacterial exposure, we wanted to further annotate the tumor selective mechanism of bacterial colonization. We hypothesized that indiscriminately seeded bacteria are more likely to survive in the immunosuppressed tumor microenvironment compared to immunocompetent tissues. To quantify bacterial presence, we performed biodistribution studies on immunocompetent and immunosuppressed tissues of survival study KPC mice at end point. In general, primary PDAC tissue, keratoacanthomas, and visible metastatic lesions contained more bacteria compared to healthy, non-target tissues (Fig 3E). Lungs and liver tissue from treated mice exhibited evidence of both gross and microscopic metastases, as identified by CK19 staining (Supp Fig 3C). Co-staining for both CK19 and O6, a serotype marker for Nissle 1917 (*22*), revealed co-localization of bacteria in CK19+ regions of lung and liver tissues, as well as the primary PDAC tumors (Fig 3F).

Furthermore, co-staining for both O6 and CC3 also revealed co-localization of bacteria and apoptotic cells, confirming cytotoxic effect of Theta production in the vicinity of tumor cells (Supp Fig 3D). Together, these data demonstrate that therapeutic bacteria can colonize both primary and metastatic tumor sites, though bacteria in all target tissues eventually decrease in density and payload production.

While treatment with Nis-Theta conferred a therapeutic benefit, its effects on normal tissues were still unknown. To evaluate this, we examined both tumors and non-target tissues for indications of cellular damage potentially caused by bacterial treatment. In Nis-Theta treated mice, the only tissue type that exhibited evidence of cell death was tumor tissue, including both primary PDAC and liver metastases (Fig 3G, Supp Fig 4). Nis-Theta treated tumors possessed significantly more apoptotic cells than both Nis-GFP treated tumors and neighboring, nontargeted tissues. Taken together, these data indicate that engineered bacteria can be safely administered to generate significant, therapeutic benefit in murine PDAC models.

### Probiotic bacteria recruit anti-tumor immune cell populations to the PDAC microenvironment

Regardless of payload, bacteria alone can induce modest recruitment of anti-tumor immune cells to previously immunologically “cold” tumors (*28, 29*). In this study, treated tumors regularly presented with liquid cores, suggestive of significant immune infiltration.

Overall, we observed no significant change in the total immune cell (CD45+) population across treatment types (Fig 4A). However, Nis-Theta treated tumors exhibited increases in helper T-cell (CD4+) and cytotoxic T-cell (CD8+) populations (Fig 4B-D). Additionally, Nis-Theta treatment also demonstrated increased cytotoxic T-cell activation (Granzyme B+) (Fig 4E). Nis-GFP treated tumors only presented significant increases in total T-cells (CD3+) (Fig 4B), suggesting an adjuvant-like contribution of Theta toxin. In contrast to the influx of other T-cell populations, we did not observe any changes in regulatory T-cells (FOXP3+) (Fig 5A).

**Fig. 4.**
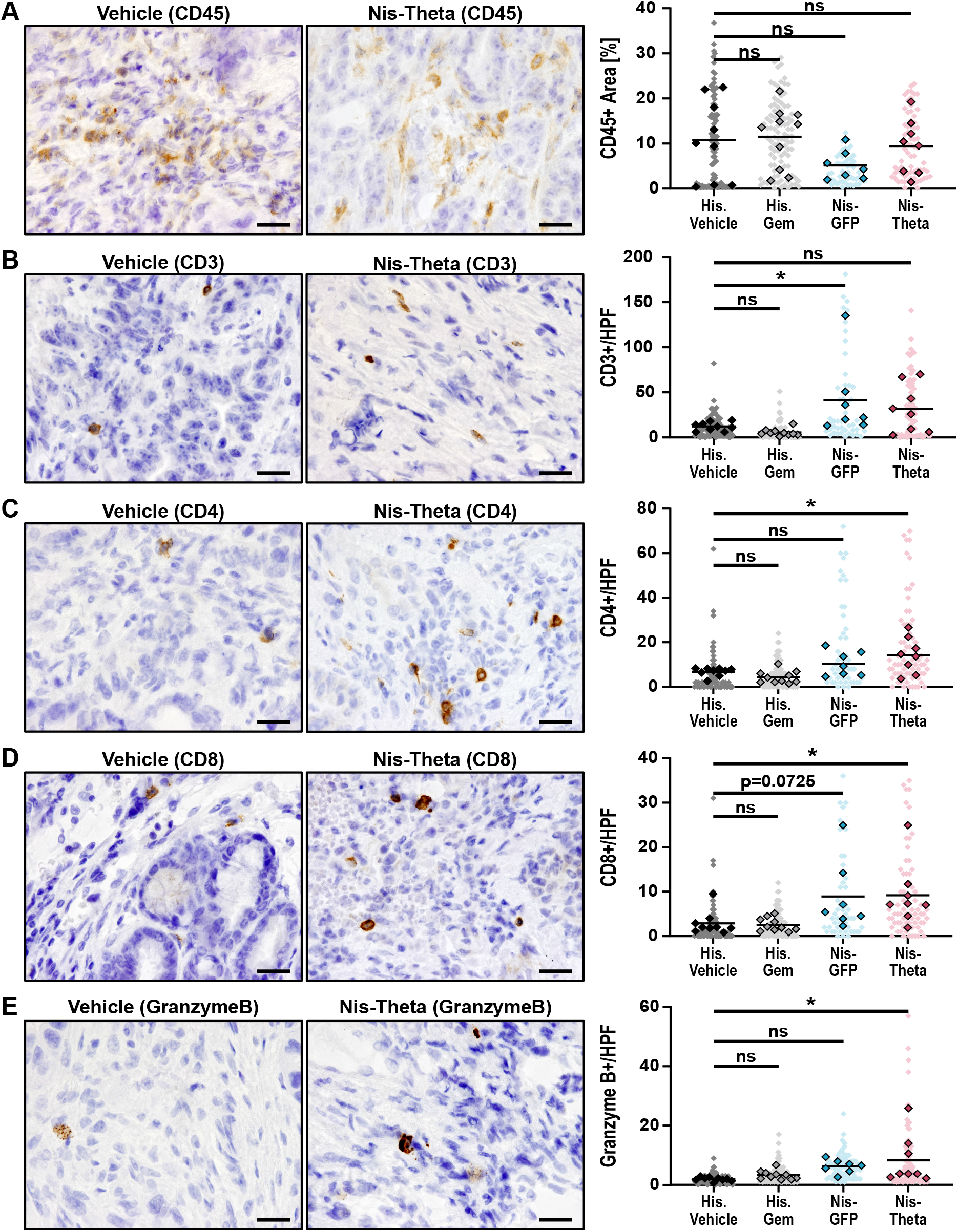
Nis-Theta treated tumors exhibit an increase in potentially anti-tumor lymphoid populations. (**A**) Pan-immune cell lineage marker, CD45. (**B**) Pan-T cell marker, CD3. (**C**) Helper-T cell marker, CD4. (**D**) Cytotoxic-T cell marker, CD8. (**E**) Granzyme B activation marker for T-cells. Representative images of vehicle and Nis-Theta treated tumors shown at 100x magnification, scale bar = 20 μm. Quantification of IHC staining based on 10-12 fields of view at 40x magnification (light shade) as either total pixel area (% Area) or discrete cell count per high power field (HPF), averaged per tumor (solid shade). Bars indicate mean ± SD. One-way ANOVA (Dunnett’s correction). \**P*<0.05 and ns, not significant.

**Fig. 5.**
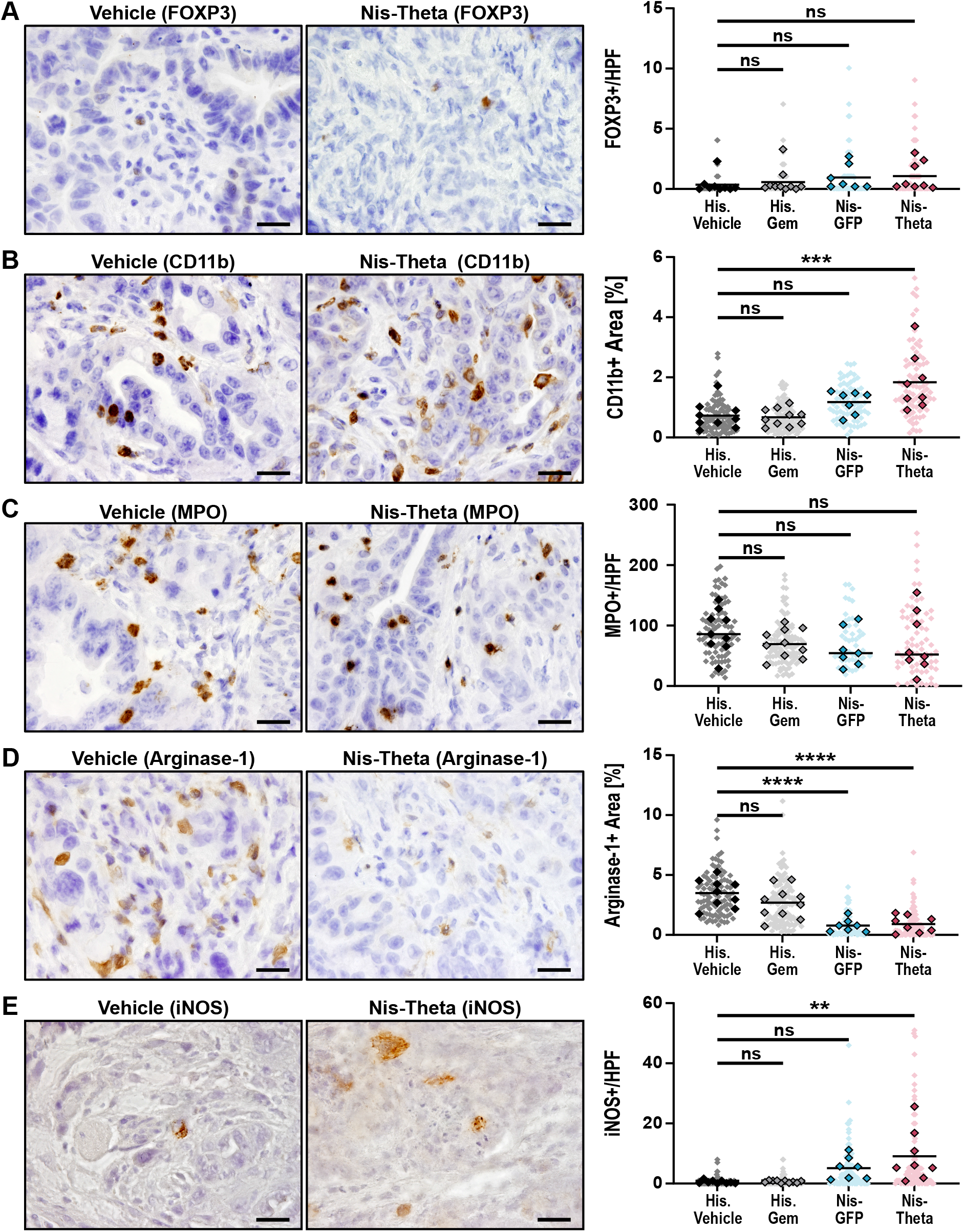
Bacteria treated tumors exhibit decreased immunosuppressive phenotypes, despite an increase in overall myeloid populations. (**A**) Regulatory-T cell marker, FOXP3. (**B**) Pan-myeloid cell lineage marker, CD11b. (**C**) Activated neutrophil marker, MPO. (**D**) M2a macrophage and myeloid derived suppressor cells (MDSC) marker, Arginase-1. (**E**) M1a macrophage marker, iNOS. Representative images of vehicle and Nis-Theta treated tumors shown at 100x magnification, scale bar = 20 μm. Quantification of IHC staining based on 10-12 fields of view at 40x magnification (light shade) as either total pixel area (% Area) or discrete cell count per high power field (HPF), averaged per tumor (solid shade). Bars indicate mean ± SD. One-way ANOVA (Dunnett’s correction). \*\**P*<0.01, \*\*\**P*<0.001, \*\*\*\**P*<0.0001, and ns, not significant.

We did observe a significant increase in CD11b+ signal for overall myeloid lineage cells (Fig 5B). Surprisingly, while we anticipated that the increase in overall myeloid populations was evidence of an innate immune response, there was no observable difference in neutrophil activation (myeloperoxidase, MPO) (Fig 5C), despite the bacterial presence. Despite this increase in potential immunosuppressive populations, markers of immunosuppressive phenotype (Arginase-1+) were significantly downregulated (Fig 5D) while iNOS, a marker of M1 macrophage activation, was upregulated (Fig 5E). Together, these data suggest that despite an overall increase in myeloid cells, bacteria-treated tumors exhibit some indications of a less immunosuppressive microenvironment.

## DISCUSSION

PDAC is a complicated and enigmatic disease. Despite its aggressive nature, it often is not diagnosed until it has spread to other vital organs. Additionally, PDAC actively suppresses angiogenesis, not only limiting access to oxygen and nutrients, but also creating a biophysical hurdle for small molecule chemotherapeutic delivery. The limitations associated with surgical intervention and intravenous therapeutics highlight the critical need for alternative methods of drug delivery for PDAC.

Here, we demonstrated the potential utility of a “living drug” capable of adapting to and thriving in the harsh PDAC tumor microenvironment. We found that therapeutic strains based on the probiotic *E. coli* Nissle 1917 can synthesize cytotoxic molecules directly within the tumor parenchyma, leading to measurable tumor control with minimal impact on normal tissues. Because the relatively non-specific pore-forming method of action makes Theta toxin potentially dangerous to cells of any lineage (*14*), systemic administration of purified toxin protein would harm normal tissues. Instead, we leverage the immunosuppression of PDAC as an advantage, providing a safe harbor for therapeutic bacteria to colonize tumor tissues while allowing their clearance from normal tissues and circulation. These data serve as proof-of-principle for approaches that harness bacteria to synthesize anti-cancer agents locally in tumor tissues.

The ability of the bacteria to colonize distant tumor sites following a local injection while sparing the non-diseased tissues further demonstrates the strength of this living drug, particularly for a disease such as PDAC that is characterized by widespread metastatic dissemination. Nearly all human PDAC cases are biopsied, either through interventional endoscopy or interventional radiology approaches. The application of a living drug during standard diagnostic procedures offers the opportunity to immediately initiate anti-cancer intervention and potentially stymie metastatic dissemination at an early stage, with little additional risk to the patient. Incorporation of additional synthetic biology tools could further enable improved tumor diagnostics (*30, 31*) and delivery of anti-cancer biological agents such therapeutic nanobodies that are otherwise considered potentially systemically toxic (*28, 29, 32*). Our data demonstrate an approach that could easily align with current standard-of-care procedures while offering the potential of prolonged tumor colonization and treatment from a single dose of living drug.

The characteristic local immunosuppressive nature of PDAC tumors continues to challenge the field of immunotherapy; to date only a few percent of patients qualify for treatment with the clinically-approved immunologic agents (*33–36*). Although evidence suggests that combination with chemotherapy enhances the effectiveness of immunotherapy (*37*), chemotherapy induced neutropenia (*38*) increases susceptibility to opportunistic bacterial infections (*39–44*), posing serious risks to patients. While the infection mechanics of *Salmonella*, *Listeria*, and *Clostridium* make them attractive candidates for effective tumoral colonization, the attenuation methods to make them safe often negate the natural tumor-targeting advantages and require additional interventions to restore tumor targeting (*45–54*). In contrast, the inherent immunogenicity of a non-native, but naturally non-pathogenic probiotic bacteria such as Nissle 1917 presents the opportunity to induce changes to the local immune microenvironment without risking more serious disease.

Here, the changes observed in the anti-tumor immune cell populations of bacteria-treated tumors may account for a portion of the tumor response. Additionally, the robust reduction in immunosuppressive phenotypes is perhaps a more significant finding in the context of PDAC specifically. Future work will investigate whether this effect may sensitize bacteria-treated tumors to combination immunotherapy. While bacteria have previously demonstrated efficacy as vectors for immune stimulating agents such as tetanus toxoid (*53*), IL-2 (*7, 8*), CD47 (*28*), PD1 and CTLA4 (*29*) in a pre-clinical setting, generation of a single bacterial vector that both directly kills tumor cells and also delivers immunotherapeutic agents could further potentiate anti-tumor effects while still being packaged as a single therapeutic agent. The data presented here provide strong rationale for the exploration of living bacterial therapy as a candidate therapeutic avenue for PDAC.

## MATERIALS AND METHODS

### Study Design

The objective of the study was to develop a strategy for tumor-specific delivery of a cytotoxic molecule using an engineered, probiotic strain of *E. coli* (Nissle 1917) in a clinically predictive animal model (KPC) of pancreatic ductal adenocarcinoma (PDAC). 2D cell lines and 3D tumor explants were first used to screen a selection of bacterially produced cytotoxins for maximum toxicity against PDAC tissues, ultimately identifying Theta toxin as a top candidate. Live, AHL-inducible bacteria producing either non-toxic GFP or Theta toxin were then administered to autochthonous PDAC tumors via direct intratumoral injection. Tumor growth and bacterial presence were monitored throughout the study by ultrasound scans and bioluminescence imaging, respectively. Overall survival and tumor doubling time were the primary metrics for therapeutic efficacy, compared to historical vehicle and chemotherapy (gemcitabine) treated cohorts. Bacterial colonization and spread were examined using immunohistochemistry and quantified by biodistribution on tumor and peripheral, non-target tissues. Systemic toxicity was tested through immunohistochemistry of tumors and peripheral, non-target tissues for markers of apoptosis. Changes in tumor microenvironment immune populations were quantified via immunohistochemistry and compared to historical treated cohorts.

### Animal Models

All animal research experiments were approved by the Columbia University Irving Medical Center (CUIMC) Institutional Animal Care and Use Committee. Mouse colonies were bred and maintained with standard mouse chow and water, *ad libitum*, under a standard 12hr light/12hr dark cycle. KPC (*Kras*^LSL.G12D/+^; *P53*^LSL.R172H/+^; *Pdx1-Cre*), KC (*Kras*^LSL.G12D/+^; *Pdx1-Cre*), and PC (*P53*^LSL.R172H/+^; *Pdx1-Cre*) mice (*15, 55*) were generated in the Olive Laboratory by crossing the described alleles. Mouse genotypes were determined using real time PCR with specific probes designed for each gene (Transnetyx; Cordova, TN). Study endpoint was determined by a health scoring system approved by IACUC.

### Bacterial Strains & Plasmids

Toxin plasmids were constructed using Gibson assembly or standard restriction enzyme-mediated cloning methods. *Salmonella typhimurium* strains (parental-ELH1301, plasmid base-pTH05, antibiotic resistance-kanamycin) previously described (*16*) were used for toxin lysate production experiments. All live bacterial experiments were performed using *Escherichia coli* (subspecies Nissle 1917), with an integrated *luxCDABE* cassette with erythromycin marker (Nislux strain). Inducible payload plasmids were derived from a p246 backbone with an integrated region from pTH05 plasmid containing *luxR, luxR* promoter, and therapeutic sequences under the control of *luxI* promoter. Payload production was induced with 10 μM acylhomoserine lactone (AHL). The AxeTxe stabilizing system was introduced to minimize plasmid loss *in vivo*. Payload plasmids contained a spectinomycin marker. Therefore, Nislux with therapeutic plasmid are both erythromycin (250 μg/mL) and spectinomycin (100 μg/mL) resistant. A detailed table of toxin sequences are provided (Table S1).

### Bacterial Toxin Lysate Preparation

Toxin producing *S. typhimurium* bacteria strains were cultured overnight in LB media containing appropriate antibiotics. Cultures were diluted 1:1000 into LB media with antibiotics and 10 μM AHL and grown for 7 hours. All cultures were normalized with additional LB to the culture with the lowest OD600 density, then 5 mL were collected and spun down. Supernatant was frozen separately. Each bacterial cell pellet was then resuspended in 2 mL PBS and subjected to 7-10 rounds of freeze-thawing in liquid nitrogen and 37°C water bath, respectively. Lysate solutions were aliquoted and stored at -20°C until use.

### Monolayer Cell Lines & *in vitro* Experiments

All human cell lines were obtained from ATCC and tested negatively for mycoplasma infection. All murine cell lines were previously derived from KPC tumors and maintained in-house. All cells are maintained under standard conditions (37°C and 5% CO_2_) and cultured in DMEM (Life Technologies, 12430-054) supplemented with 1% penicillin and streptomycin (Corning, 30-003-CI), 10% fetal bovine serum (Life Technologies, 10438-034), and 1% MEM non-essential amino acids (Fischer Scientific, 11-140-050).

Cells were seeded at 2,500 cells/well in 96-well plates and allowed to adhere overnight. The next day, media was aspirated and replaced with 100μL standard media supplemented with 10% bacteria lysate stock. After 44 hours, 10 μL of Alamar Blue (BioRad, BUF012B) was added and incubated at 37°C, 5% CO_2_ for 4 hours, then fluorescence (ex. 560 nm, em. 590 nm) was measured on a Promega multimode microplate reader. Background levels were subtracted from raw results, then further normalized as a percentage of PBS treated samples. This assay was completed three times with four technical replicates per biological replicate.

GFP and β Hemolysin (βHemo) were selected as “ineffective” payloads for comparison to Hemolysin E (HlyE), Heat Stable Enterotoxin (HSEnt), and Theta as “effective” payloads against PDAC cell lines. Magainin, a top candidate, was removed from consideration due to documented poor effectiveness on stromal cell types (*56, 57*).

### Tumor Explants & *ex vivo* Experiments

Murine tumors were collected following humane euthanasia and trimmed of healthy pancreas tissue in a sterile petri dish. Tissues were processed into explant slices and cultured as previously described (*3, 17*). Explants were treated with murine explant media supplemented with 10% bacterial toxin lysate, replaced daily for up to 3 days. For live bacteria co-cultures, bacteria strains were grown in LB media containing appropriate antibiotics for 6 hours, then spun down and washed 3 times with sterile PBS. Bacteria were concentrated to 5×10^8^ CFU/mL and 10 μL were added to each well (10^6^ total bacteria) on day 0 only. The media was changed daily, supplemented with 10 μM AHL and critically, no penicillin/streptomycin. Explants were fixed in 4% paraformaldehyde for 2 hours at 4°C, then transferred to 70% ethanol and paraffin embedded for long term storage.

### Bacterial Administration for Interventional *in vivo* Study

KPC mice were monitored by manual palpation for tumor development and presence of a tumor was confirmed via ultrasound. Accessible tumors (tail or body of pancreas) with dimensions smaller than 4-6 mm were enrolled and randomized into study arms. *Post hoc* analysis determined no significant enrichment for sex in any arm of the studies was observed. The mice were first started on a 7-10 day pre-treatment course of antibiotics. Ampicillin (1 g/L) and neomycin (0.5 g/L) were added to the drinking water and changed every three days until tumors reached dimensions between 4-6 mm. The mice were removed from antibiotics and provided with fresh water 6 hours prior to bacterial treatment.

Bacterial strains were grown in LB media containing appropriate antibiotics for 6 hours. Bacteria were spun down and washed 3 times with sterile PBS. 10x AHL was added to bacteria suspension before injection into mice. Ultrasound guided intratumoral injections of bacteria (Movie S1) were performed at a concentration of 5 x 10^8^ CFU/mL in PBS, with a total volume of 20 μL followed by a 10 μL Matrigel plug (10^7^ total bacteria injected). While on study, mice received 500μM AHL in drinking water, changed every three days.

### Monitoring Tumor and Bacterial Dynamics *in vivo*

Approximately every 3 days, on-study mice were scanned for tumor volume via ultrasound as previously described (*25*) and bacterial dynamics via an *in vivo* imaging system (AMI-HTX Optical Imaging System, Spectral Imaging) for quantification. Qualitative (non-quantification) luminescence imaging was performed with PerkinElmer, Inc. *In vivo* Imaging System (IVIS). In preparation for both scans, the abdomens of the mice were shaved, and the remaining stubble removed with depilatory cream (Nair).

Two scans for constitutive luminescent signal from the integrated *luxCDABE* cassette were performed for 45 second and 2 minutes respectively at both MP6 (supine position) and MP3 (laying on right flank). Luminescent signal was quantified and analyzed with Aura Imaging Software (Spectral Imaging), where background radiance flux (photons/s) from identical sized ROIs was subtracted from the tumor region flux. Representative images were scaled to a single, consistent threshold for presentation.

### Biodistribution

At survival end point, all tissues were harvested. Tumors, spleen, lung, liver, and metastatic tissue and/or keratoacanthomas when applicable, were harvested. Half of each select organ (tumor, spleen, lung, liver, and metastatic site and/or keratoacanthomas where applicable) were collected into sterile PBS, while the other half and all other tissues were fixed in 10% formalin overnight. Fresh collected tissues were weighed and homogenized using Polytron PT 1300D homogenizer. Homogenates were serially diluted and spot plated, 5 μL, on LB-agar plates containing appropriate antibiotics at 37°C overnight. Colonies were counted and computed as CFU/g tissue (limit of-detection 10^3^ CFU/g).

### Immunohistochemistry

Formalin-fixed, paraffin-embedded tissues were sectioned in 4 μm slides and rehydrated using a Leica XL ST5010 autostainer. Heat-activated antigen retrieval was performed followed by incubation in 3% hydrogen peroxide to quench endogenous peroxidases prior to incubation with primary antibodies (Table S2). Secondary antibody incubation and developed with either ImmPACT DAB Peroxidase (HRP) Substrate (Vector Laboratories, SK-4105) or VIP Peroxidase (HRP) Substrate (Vector Laboratories, SK-4600), followed by hematoxylin counterstain, dehydration, and coverslip mounting. Quantitative analyses of IHC images were performed using FIJI (*58*).

### Western Blot

Bacteria were lysed in 500 μL Bugbuster buffer (Millipore Sigma, 70584) supplemented with 0.5 μL rLysozyme (Millipore Sigma, 71110) and 0.5 μL Benzonase endonuclease (Millipore Sigma, 101697). Supernatants and lysates were concentrated using Merck Millipore Ultracel -10K centrifugal filters (UFC501024). 20 µl of lysate or supernatant were loaded on sodium dodecyl sulfate-polyacrylamide gels (BioRad, 4561093) for gel electrophoresis. Blotting was performed using TurboBlot system (BioRad, 10026938). After 1 hour blocking in 5% bovine serum albumin (BSA) in TBS+0.1% Tween20, protein membranes were incubated with PFO antibody (1:1,000; MyBioSource, MBS7049795) at 4 °C overnight. Secondary antibodies were applied for 1 hour at room temperature (1:5,000; CST, #7074) before detecting protein bands at the BioRad ChemiDoc MP Imaging System using chemiluminescence (BioRad, 170-5061).

### Statistical Analysis

Analyses were performed using GraphPad Prism v9.5.1. Comparison across multiple variables used two-way ANOVA with either Dunnett’s (cell line toxin selection) or Bonferroni’s (tissue apoptosis) corrections. Comparison of multiple data groups to one reference group (*ex vivo* explants, tumor doubling time, immune populations) were tested using one-way ANOVA with Dunnett’s correction. Group-to-group differences of bacterial colonization data were determined using Mann-Whitney t-test (non-parametric test). For all analyses, *P*<0.05 was considered statistically significant. \**P*<0.05, *\*\*P*<0.01, \*\*\**P*<0.001, and \*\*\*\**P*<0.0001.

### List of Supplementary Materials

Figs. S1 to S4

Table S1 to S2

Movie S1

## Acknowledgments

We thank and acknowledge the contributions of the Mouse Hospital, Oncology Precision Therapeutics and Imaging Core (OPTIC), and Molecular Pathology Shared Resources (MSRP) of Columbia University. We also thank Dr. Martina Pavlicova for her insight regarding statistical analyses.

## Funding

National Institutes of Health grant R01CA215607 (KPO)

National Institutes of Health grant U01CA274312 (KPO)

Lustgarten Foundational Clinical Translational Program Award (KPO)

National Institutes of Health grant R01EB029750 (TD)

National Institutes of Health grant F31CA250443 (ARDF)

Deutsche Forschungsgemeinschaft 439440500 (MCH)

National Institutes of Health grant F99CA253756 (TH)

Honjo International Foundation Scholarship (TH)

Herbert Irving Comprehensive Cancer Center Cancer Center Support Grant P30CA013696 (Columbia University)

## Author contributions

Conceptualization: ARDF, TD, KPO

Resources: SAS, TH, CFP, ERB, TD, KPO

Data curation: ARDF, TD, KPO

Formal analysis: ARDF, MCH, CFP

Supervision: SAS, MAB, TD, KPO

Funding acquisition: ARDF, TD, KPO

Validation: ARDF, TD, KPO

Investigation: ARDF, SAS, MCH, TD, KPO

Visualization: ARDF, MCH, MAB, TD, KPO

Methodology: ARDF, SAS, TH, MCH, CFP, ERB, TD, KPO

Writing – original draft: ARDF

Project administration: ARDF, TD, KPO

Writing – review and editing: ARDF, MCH, MAB, TD, KPO

## Data and materials availability

All data generated in this study are available upon request from the corresponding author.

## References and Notes

1. R. L. Siegel, K. D. Miller, N. S. Wagle, A. Jemal, Cancer statistics, 2023. CA Cancer J Clin 73, 17–48 (2023).

2. W. Park, A. Chawla, E. M. O’Reilly, Pancreatic Cancer: A Review. Jama 326, 851–862 (2021).

3. M. C. Hasselluhn, A. R. Decker-Farrell, L. Vlahos, D. H. Thomas, A. Curiel-Garcia, H. C. Maurer, U. N. Wasko, L. Tomassoni, S. A. Sastra, C. F. Palermo, T. C. Dalton, A. Ma, F. Li, E. J. Tolosa, H. Hibshoosh, M. E. Fernandez-Zapico, A. Muir, A. Califano, K. P. Olive, “Tumor Explants Elucidate a Cascade of Paracrine SHH, WNT, and VEGF Signals Driving Pancreatic Cancer Angiosuppression”. bioRxiv, 2023.2003.2002.529724 (2023).

4. K. P. Olive, M. A. Jacobetz, C. J. Davidson, A. Gopinathan, D. McIntyre, D. Honess, B. Madhu, M. A. Goldgraben, M. E. Caldwell, D. Allard, K. K. Frese, G. Denicola, C. Feig, C. Combs, S. P. Winter, H. Ireland-Zecchini, S. Reichelt, W. J. Howat, A. Chang, M. Dhara, L. Wang, F. Ruckert, R. Grutzmann, C. Pilarsky, K. Izeradjene, S. R. Hingorani, P. Huang, S. E. Davies, W. Plunkett, M. Egorin, R. H. Hruban, N. Whitebread, K. McGovern, J. Adams, C. Iacobuzio-Donahue, J. Griffiths, D. A. Tuveson, Inhibition of Hedgehog signaling enhances delivery of chemotherapy in a mouse model of pancreatic cancer. Science 324, 1457–1461 (2009).

5. A. Di Federico, M. Mosca, R. Pagani, R. Carloni, G. Frega, A. De Giglio, A. Rizzo, D. Ricci, S. Tavolari, M. Di Marco, A. Palloni, G. Brandi, Immunotherapy in Pancreatic Cancer: Why Do We Keep Failing? A Focus on Tumor Immune Microenvironment, Predictive Biomarkers and Treatment Outcomes. Cancers (Basel) 14, (2022).

6. R. Hassan, M. Ho, Mesothelin targeted cancer immunotherapy. Eur J Cancer 44, 46–53 (2008).

7. G. Batist, Park J.Y., Drees J., Kangas T., Saltzman D., Orally Administered Multiple Dose Saltikva (Salmonella-IL2) in Conjunction with Folfirinox in a Patient with Stage IV Pancreatic Cancer: A Case Report. Clin Oncol Case Rep 3, (2020).

8. T. J. Gniadek, L. Augustin, J. Schottel, A. Leonard, D. Saltzman, E. Greeno, G. Batist, A Phase I, Dose Escalation, Single Dose Trial of Oral Attenuated Salmonella typhimurium Containing Human IL-2 in Patients With Metastatic Gastrointestinal Cancers. J Immunother 43, 217–221 (2020).

9. F. P. Canale, C. Basso, G. Antonini, M. Perotti, N. Li, A. Sokolovska, J. Neumann, M. J. James, S. Geiger, W. Jin, J.-P. Theurillat, K. A. West, D. S. Leventhal, J. M. Lora, F. Sallusto, R. Geiger, Metabolic modulation of tumours with engineered bacteria for immunotherapy. Nature 598, 662–666 (2021).

10. C. R. Gurbatri, N. Arpaia, T. Danino, Engineering bacteria as interactive cancer therapies. Science 378, 858–864 (2022).

11. Y. E. Chen, D. Bousbaine, A. Veinbachs, K. Atabakhsh, A. Dimas, V. K. Yu, A. Zhao, N. J. Enright, K. Nagashima, Y. Belkaid, M. A. Fischbach, Engineered skin bacteria induce antitumor T cell responses against melanoma. Science 380, 203–210 (2023).

12. T. M. Savage, R. L. Vincent, S. S. Rae, L. H. Huang, A. Ahn, K. Pu, F. Li, K. de los Santos-Alexis, C. Coker, T. Danino, N. Arpaia, Chemokines expressed by engineered bacteria recruit and orchestrate antitumor immunity. Science Advances 9, eadc9436 (2023).

13. S. Jiang, G. Redelman-Sidi, BCG in Bladder Cancer Immunotherapy. Cancers (Basel) 14, (2022).

14. S. Verherstraeten, E. Goossens, B. Valgaeren, B. Pardon, L. Timbermont, F. Haesebrouck, R. Ducatelle, P. Deprez, K. R. Wade, R. Tweten, F. Van Immerseel, Perfringolysin O: The Underrated Clostridium perfringens Toxin? Toxins (Basel) 7, 1702–1721 (2015).

15. S. R. Hingorani, L. Wang, A. S. Multani, C. Combs, T. B. Deramaudt, R. H. Hruban, A. K. Rustgi, S. Chang, D. A. Tuveson, Trp53R172H and KrasG12D cooperate to promote chromosomal instability and widely metastatic pancreatic ductal adenocarcinoma in mice. Cancer Cell 7, 469–483 (2005).

16. T. Harimoto, D. Deb, T. Danino, A rapid screening platform to coculture bacteria within tumor spheroids. Nat Protoc 17, 2216–2239 (2022).

17. A. R. Decker-Farrell, A. Ma, F. Li, A. Muir, K. P. Olive, Generation and ex vivo culture of murine and human pancreatic ductal adenocarcinoma tissue slice explants. STAR Protoc 4, 102711 (2023).

18. J. Stritzker, S. Weibel, P. J. Hill, T. A. Oelschlaeger, W. Goebel, A. A. Szalay, Tumor-specific colonization, tissue distribution, and gene induction by probiotic Escherichia coli Nissle 1917 in live mice. Int J Med Microbiol 297, 151–162 (2007).

19. T. Danino, A. Prindle, G. A. Kwong, M. Skalak, H. Li, K. Allen, J. Hasty, S. N. Bhatia, Programmable probiotics for detection of cancer in urine. Sci Transl Med 7, 289ra284 (2015).

20. R. Li, L. Helbig, J. Fu, X. Bian, J. Herrmann, M. Baumann, A. F. Stewart, R. Muller, A. Li, D. Zips, Y. Zhang, Expressing cytotoxic compounds in Escherichia coli Nissle 1917 for tumor-targeting therapy. Res Microbiol 170, 74–79 (2019).

21. I. Gentschev, I. Petrov, M. Ye, L. Kafuri Cifuentes, R. Toews, A. Cecil, T. A. Oelschaeger, A. A. Szalay, Tumor Colonization and Therapy by Escherichia coli Nissle 1917 Strain in Syngeneic Tumor-Bearing Mice Is Strongly Affected by the Gut Microbiome. Cancers (Basel) 14, (2022).

22. L. Grozdanov, C. Raasch, J. Schulze, U. Sonnenborn, G. Gottschalk, J. Hacker, U. Dobrindt, Analysis of the genome structure of the nonpathogenic probiotic Escherichia coli strain Nissle 1917. J Bacteriol 186, 5432–5441 (2004).

23. U. Sonnenborn, J. Schulze, The non-pathogenic Escherichia coli strain Nissle 1917 – features of a versatile probiotic. Microbial Ecology in Health and Disease 21, 122–158 (2009).

24. A. J. H. Fedorec, T. Ozdemir, A. Doshi, Y. K. Ho, L. Rosa, J. Rutter, O. Velazquez, V. B. Pinheiro, T. Danino, C. P. Barnes, Two New Plasmid Post-segregational Killing Mechanisms for the Implementation of Synthetic Gene Networks in Escherichia coli. iScience 14, 323–334 (2019).

25. S. A. Sastra, K. P. Olive, Quantification of murine pancreatic tumors by high-resolution ultrasound. Methods Mol Biol 980, 249–266 (2013).

26. J. A. Eberle-Singh, I. Sagalovskiy, H. C. Maurer, S. A. Sastra, C. F. Palermo, A. R. Decker, M. J. Kim, J. Sheedy, A. Mollin, L. Cao, J. Hu, A. Branstrom, M. Weetall, K. P. Olive, Effective Delivery of a Microtubule Polymerization Inhibitor Synergizes with Standard Regimens in Models of Pancreatic Ductal Adenocarcinoma. Clin Cancer Res 25, 5548–5560 (2019).

27. N. M. Gades, A. Ohash, L. D. Mills, M. A. Rowley, K. S. Predmore, R. J. Marler, F. J. Couch, Spontaneous vulvar papillomas in a colony of mice used for pancreatic cancer research. Comp Med 58, 271–275 (2008).

28. S. Chowdhury, S. Castro, C. Coker, T. E. Hinchliffe, N. Arpaia, T. Danino, Programmable bacteria induce durable tumor regression and systemic antitumor immunity. Nat Med 25, 1057–1063 (2019).

29. C. R. Gurbatri, I. Lia, R. Vincent, C. Coker, S. Castro, P. M. Treuting, T. E. Hinchliffe, N. Arpaia, T. Danino, Engineered probiotics for local tumor delivery of checkpoint blockade nanobodies. Sci Transl Med 12, (2020).

30. A. A. Ordonez, F. Saupe, C. A. Kasper, M. L. Turner, S. Parveen, K. Flavahan, H. Shin, D. Artemov, S. J. Ittig, S. K. Jain, Imaging Tumor-Targeting Bacteria Using 18F-Fluorodeoxysorbitol Positron Emission Tomography. J Infect Dis 228, S291–s296 (2023).

31. T. Jiang, X. Yang, G. Li, X. Zhao, T. Sun, R. Müller, H. Wang, M. Li, Y. Zhang, Bacteria-Based Live Vehicle for In Vivo Bioluminescence Imaging. Analytical Chemistry 93, 15687-15695 (2021).

32. M. H. Abedi, M. S. Yao, D. R. Mittelstein, A. Bar-Zion, M. B. Swift, A. Lee-Gosselin, P. Barturen-Larrea, M. T. Buss, M. G. Shapiro, Ultrasound-controllable engineered bacteria for cancer immunotherapy. Nature Communications 13, 1585 (2022).

33. G. J. Weiss, J. Waypa, L. Blaydorn, J. Coats, K. McGahey, A. Sangal, J. Niu, C. A. Lynch, J. H. Farley, V. Khemka, A phase Ib study of pembrolizumab plus chemotherapy in patients with advanced cancer (PembroPlus). Br J Cancer 117, 33–40 (2017).

34. T. Doi, K. Muro, H. Ishii, T. Kato, T. Tsushima, M. Takenoyama, S. Oizumi, K. Gemmoto, H. Suna, K. Enokitani, T. Kawakami, H. Nishikawa, N. Yamamoto, A Phase I Study of the Anti-CC Chemokine Receptor 4 Antibody, Mogamulizumab, in Combination with Nivolumab in Patients with Advanced or Metastatic Solid Tumors. Clin Cancer Res 25, 6614–6622 (2019).

35. M. Ghidini, A. Lampis, M. B. Mirchev, A. F. Okuducu, M. Ratti, N. Valeri, J. C. Hahne, Immune-Based Therapies and the Role of Microsatellite Instability in Pancreatic Cancer. Genes (Basel) 12, (2020).

36. C. Luchini, L. A. A. Brosens, L. D. Wood, D. Chatterjee, J. I. Shin, C. Sciammarella, G. Fiadone, G. Malleo, R. Salvia, V. Kryklyva, M. L. Piredda, L. Cheng, R. T. Lawlor, V. Adsay, A. Scarpa, Comprehensive characterisation of pancreatic ductal adenocarcinoma with microsatellite instability: histology, molecular pathology and clinical implications. Gut 70, 148–156 (2021).

37. C. Sordo-Bahamonde, S. Lorenzo-Herrero, A. P. Gonzalez-Rodriguez, A. Martínez-Pérez, J. P. Rodrigo, J. M. García-Pedrero, S. Gonzalez, Chemo-Immunotherapy: A New Trend in Cancer Treatment. Cancers (Basel) 15, (2023).

38. G. Roviello, M. Ramello, M. Catalano, A. D’Angelo, R. Conca, S. Gasperoni, L. Dreoni, R. Petrioli, A. Ianza, S. Nobili, M. Aieta, E. Mini, Association between neutropenia and survival to nab-paclitaxel and gemcitabine in patients with metastatic pancreatic cancer. Scientific Reports 10, 19281 (2020).

39. S. Bhat, S. Muthunatarajan, S. S. Mulki, K. Archana Bhat, K. H. Kotian, Bacterial Infection among Cancer Patients: Analysis of Isolates and Antibiotic Sensitivity Pattern. Int J Microbiol 2021, 8883700 (2021).

40. A. Peretz, I. B. Shlomo, O. Nitzan, L. Bonavina, P. M. Schaffer, M. Schaffer, Clostridium difficile Infection: Associations with Chemotherapy, Radiation Therapy, and Targeting Therapy Treatments. Curr Med Chem 23, 4442–4449 (2016).

41. L. M. Noriega, P. Van der Auwera, D. Daneau, F. Meunier, M. Aoun, Salmonella infections in a cancer center. Support Care Cancer 2, 116–122 (1994).

42. N. Mori, A. D. Szvalb, J. A. Adachi, J. J. Tarrand, V. E. Mulanovich, Clinical presentation and outcomes of non-typhoidal Salmonella infections in patients with cancer. BMC Infect Dis 21, 1021 (2021).

43. G. A. Rivero, H. A. Torres, K. V. Rolston, D. P. Kontoyiannis, Listeria monocytogenes infection in patients with cancer. Diagn Microbiol Infect Dis 47, 393–398 (2003).

44. P. Mook, S. J. O’Brien, I. A. Gillespie, Concurrent conditions and human listeriosis, England, 1999-2009. Emerg Infect Dis 17, 38–43 (2011).

45. M. Zhao, M. Yang, X. M. Li, P. Jiang, E. Baranov, S. Li, M. Xu, S. Penman, R. M. Hoffman, Tumor-targeting bacterial therapy with amino acid auxotrophs of GFP-expressing Salmonella typhimurium. Proc Natl Acad Sci U S A 102, 755–760 (2005).

46. R. W. Kasinskas, N. S. Forbes, Salmonella typhimurium Lacking Ribose Chemoreceptors Localize in Tumor Quiescence and Induce Apoptosis. Cancer Research 67, 3201–3209 (2007).

47. W. Quispe-Tintaya, D. Chandra, A. Jahangir, M. Harris, A. Casadevall, E. Dadachova, C. Gravekamp, Nontoxic radioactive Listeria(at) is a highly effective therapy against metastatic pancreatic cancer. Proc Natl Acad Sci U S A 110, 8668–8673 (2013).

48. X. Xu, W. A. Hegazy, L. Guo, X. Gao, A. N. Courtney, S. Kurbanov, D. Liu, G. Tian, E. R. Manuel, D. J. Diamond, M. Hensel, L. S. Metelitsa, Effective cancer vaccine platform based on attenuated salmonella and a type III secretion system. Cancer Res 74, 6260–6270 (2014).

49. B. C. S. Dinesh Chandra, Ziqiang Yuan, Steven K Libutti,, A. B. Wade Koba, Kun Zhu, Arturo Casadevall, Ekaterina Dadachova,, C. Gravekamp, 32-Phosphorus selectively delivered by listeria to pancreatic cancer demonstrates a strong therapeutic effect. Oncotarget 8, 12 (2017).

50. R. Hassan, E. Alley, H. Kindler, S. Antonia, T. Jahan, S. Honarmand, N. Nair, C. C. Whiting, A. Enstrom, E. Lemmens, T. Tsujikawa, S. Kumar, G. Choe, A. Thomas, K. McDougall, A. L. Murphy, E. Jaffee, L. M. Coussens, D. G. Brockstedt, Clinical Response of Live-Attenuated, Listeria monocytogenes Expressing Mesothelin (CRS-207) with Chemotherapy in Patients with Malignant Pleural Mesothelioma. Clin Cancer Res 25, 5787–5798 (2019).

51. X. Feng, P. He, C. Zeng, Y. H. Li, S. K. Das, B. Li, H. F. Yang, Y. Du, Novel insights into the role of Clostridium novyi-NT related combination bacteriolytic therapy in solid tumors. Oncol Lett 21, 110 (2021).

52. L. B. Augustin, L. Milbauer, S. E. Hastings, A. S. Leonard, D. A. Saltzman, J. L. Schottel, Salmonella enterica Typhimurium engineered for nontoxic systemic colonization of autochthonous tumors. J Drug Target 29, 294–299 (2021).

53. B. C. Selvanesan, D. Chandra, W. Quispe-Tintaya, A. Jahangir, A. Patel, K. Meena, R. A. Alves Da Silva, M. Friedman, L. Gabor, O. Khouri, S. K. Libutti, Z. Yuan, J. Li, S. Siddiqui, A. Beck, L. Tesfa, W. Koba, J. Chuy, J. C. McAuliffe, R. Jafari, D. Entenberg, Y. Wang, J. Condeelis, V. DesMarais, V. Balachandran, X. Zhang, K. Lin, C. Gravekamp, Listeria delivers tetanus toxoid protein to pancreatic tumors and induces cancer cell death in mice. Sci Transl Med 14, eabc1600 (2022).

54. K. M. Dailey, J. M. Small, J. E. Pullan, S. Winfree, K. E. Vance, M. Orr, S. Mallik, K. W. Bayles, M. A. Hollingsworth, A. E. Brooks, An intravenous pancreatic cancer therapeutic: Characterization of CRISPR/Cas9n-modified Clostridium novyi-Non Toxic. PLoS One 18, e0289183 (2023).

55. S. R. Hingorani, E. F. Petricoin, A. Maitra, V. Rajapakse, C. King, M. A. Jacobetz, S. Ross, T. P. Conrads, T. D. Veenstra, B. A. Hitt, Y. Kawaguchi, D. Johann, L. A. Liotta, H. C. Crawford, M. E. Putt, T. Jacks, C. V. Wright, R. H. Hruban, A. M. Lowy, D. A. Tuveson, Preinvasive and invasive ductal pancreatic cancer and its early detection in the mouse. Cancer Cell 4, 437–450 (2003).

56. J. Lehmann, M. Retz, S. S. Sidhu, H. Suttmann, M. Sell, F. Paulsen, J. Harder, G. Unteregger, M. Stockle, Antitumor activity of the antimicrobial peptide magainin II against bladder cancer cell lines. Eur Urol 50, 141–147 (2006).

57. Y. Liscano, J. Onate-Garzon, J. P. Delgado, Peptides with Dual Antimicrobial-Anticancer Activity: Strategies to Overcome Peptide Limitations and Rational Design of Anticancer Peptides. Molecules 25, (2020).

58. J. Schindelin, I. Arganda-Carreras, E. Frise, V. Kaynig, M. Longair, T. Pietzsch, S. Preibisch, C. Rueden, S. Saalfeld, B. Schmid, J. Y. Tinevez, D. J. White, V. Hartenstein, K. Eliceiri, P. Tomancak, A. Cardona, Fiji: an open-source platform for biological-image analysis. Nat Methods 9, 676-682 (2012).

