## Supplemental Figure 1 for "“Tumor-selective treatment of metastatic pancreatic cancer with an engineered, probiotic living drug”"

### Supp Fig. 1

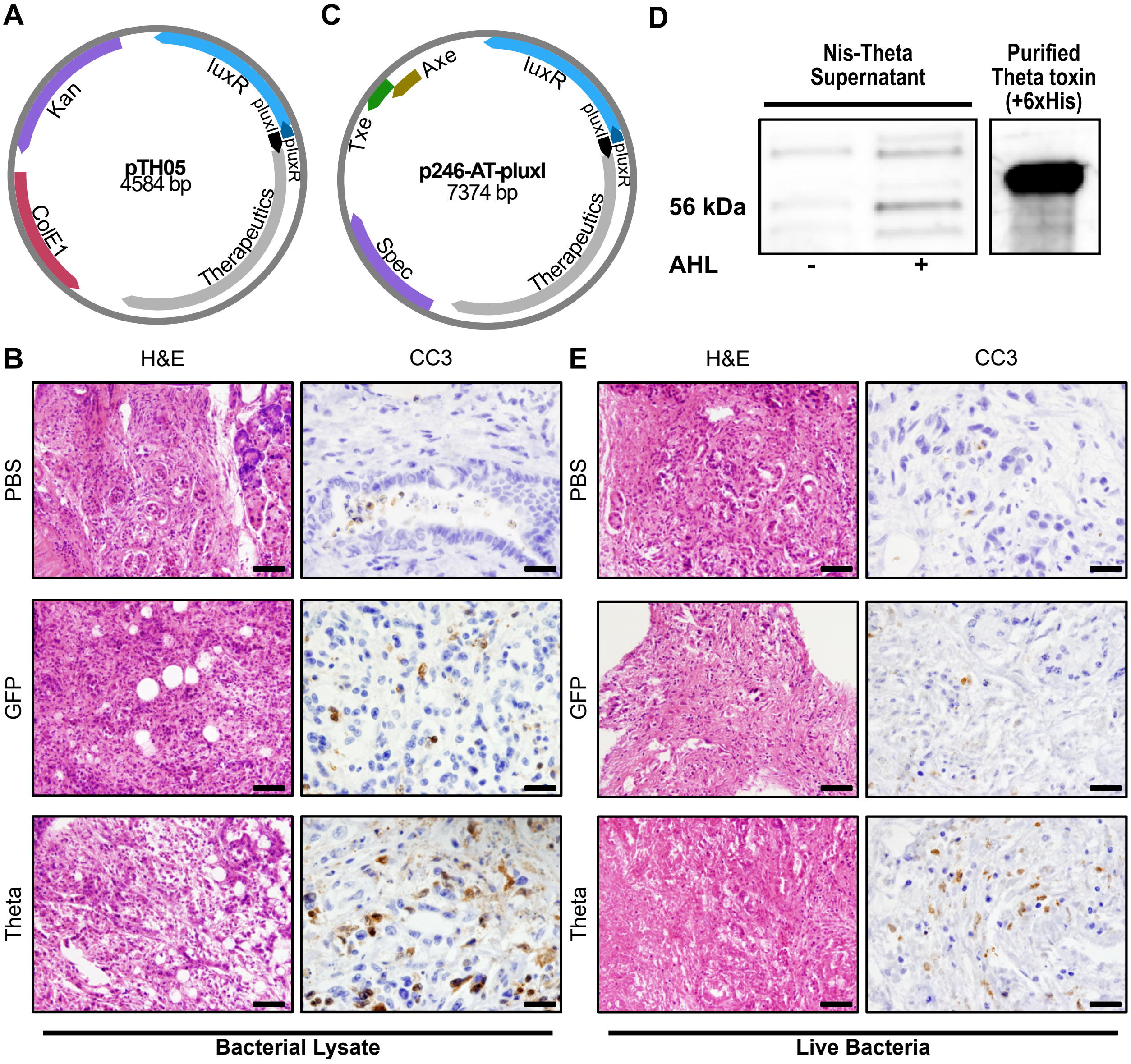


**Fig. S1. Therapeutic bacterial design and characterization.** (**A**) Map of plasmid design used in *in vitro* experiments (cell line and explant screens). pTH05, high-copy number plasmid encoding AHL induction mechanism of therapeutic production. ColE1 origin of replication and kanamycin resistance. (**B**) Representative images of explants treated for 72 hours with the indicated toxin-containing bacteria lysates for Hematoxylin & Eosin (H&E, 40x magnification, scale bar = 50 µm) and IHC for apoptosis marker CC3 (100x magnification, scale bar = 20 µm). (**C**) Map of plasmid designed used in *in vivo* experiments. p246-AT-pluxI, high-copy number plasmid encoding AHL induction mechanism of therapeutic production and AxeTxe stability mechanism. ColE1 origin of replication (not indicated) and spectinomycin resistance. (**D**) Immunoblot of Nis-Theta bacterial culture supernatant with and without AHL-induction. The immunoblot has been cropped to show relevant bands, including a purified Theta toxin construct with an additional 6x-His tag and the expected band shift. (**E**) Representative images of explants treated for 72 hours with the indicated live, toxin-producing *E. coli Nissle 1917* for 72 hours for Hematoxylin & Eosin (H&E, 40x magnification, scale bar = 50 µm) and IHC for apoptosis marker CC3 (100x magnification, scale bar = 20 µm).
