## Supplemental Figure 2 for "“Tumor-selective treatment of metastatic pancreatic cancer with an engineered, probiotic living drug”"

### Supp Fig. 2

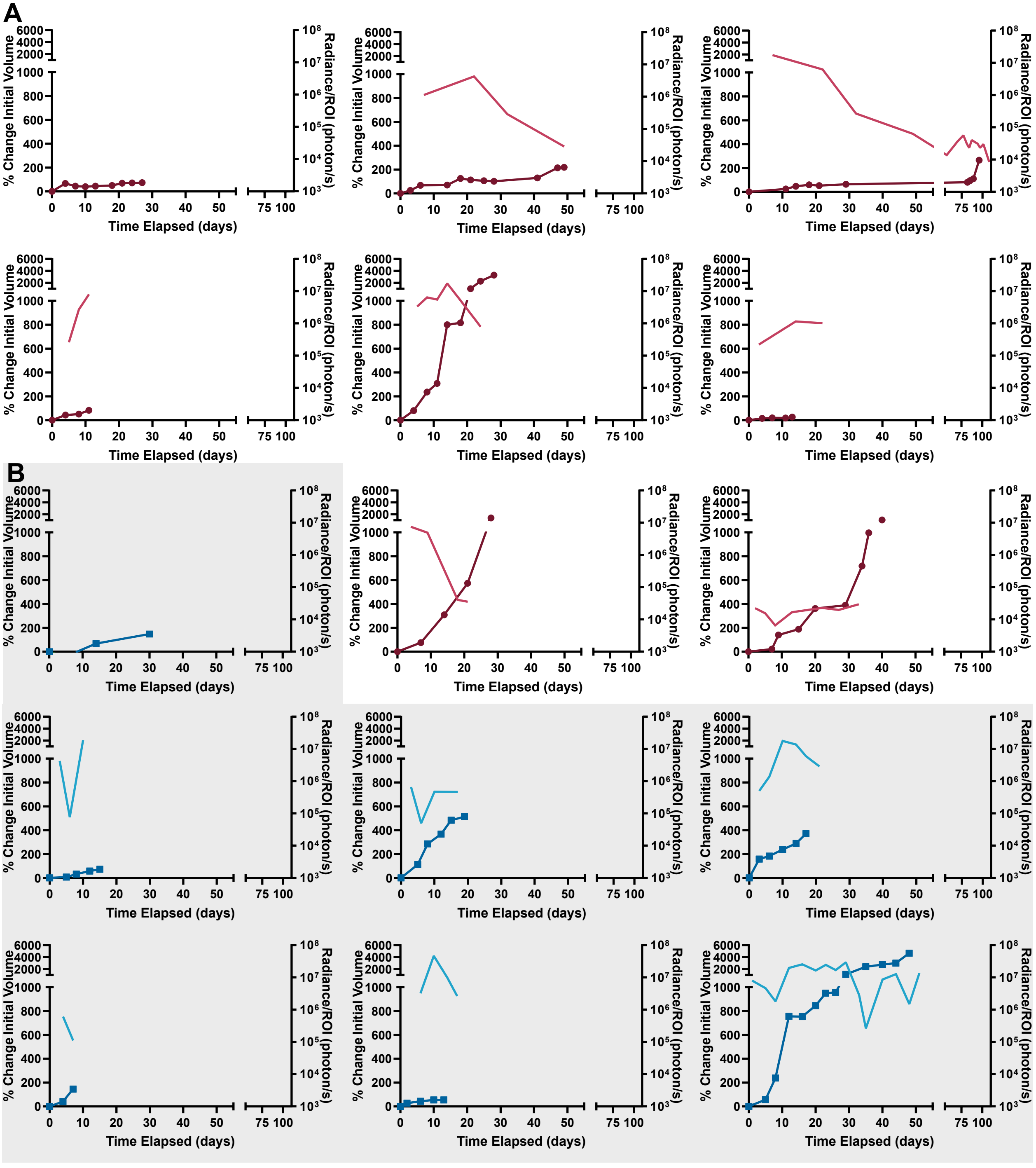


**Fig. S2.** ***In vivo*** **bacterial luminescence decreases over time on study, while tumor volume outgrow.** (**A**) Nis-Theta treated tumors. (**B**) Nis-GFP treated tumors. Individual tumor growth graces (darker color with symbols) overlayed with quantified luminescence signal (lighter color solid lines). Some tumors did not have sufficient luminescence or ultrasound scans to quantify.
