## Supplemental Figure 3 for "“Tumor-selective treatment of metastatic pancreatic cancer with an engineered, probiotic living drug”"

Supp Fig. 3

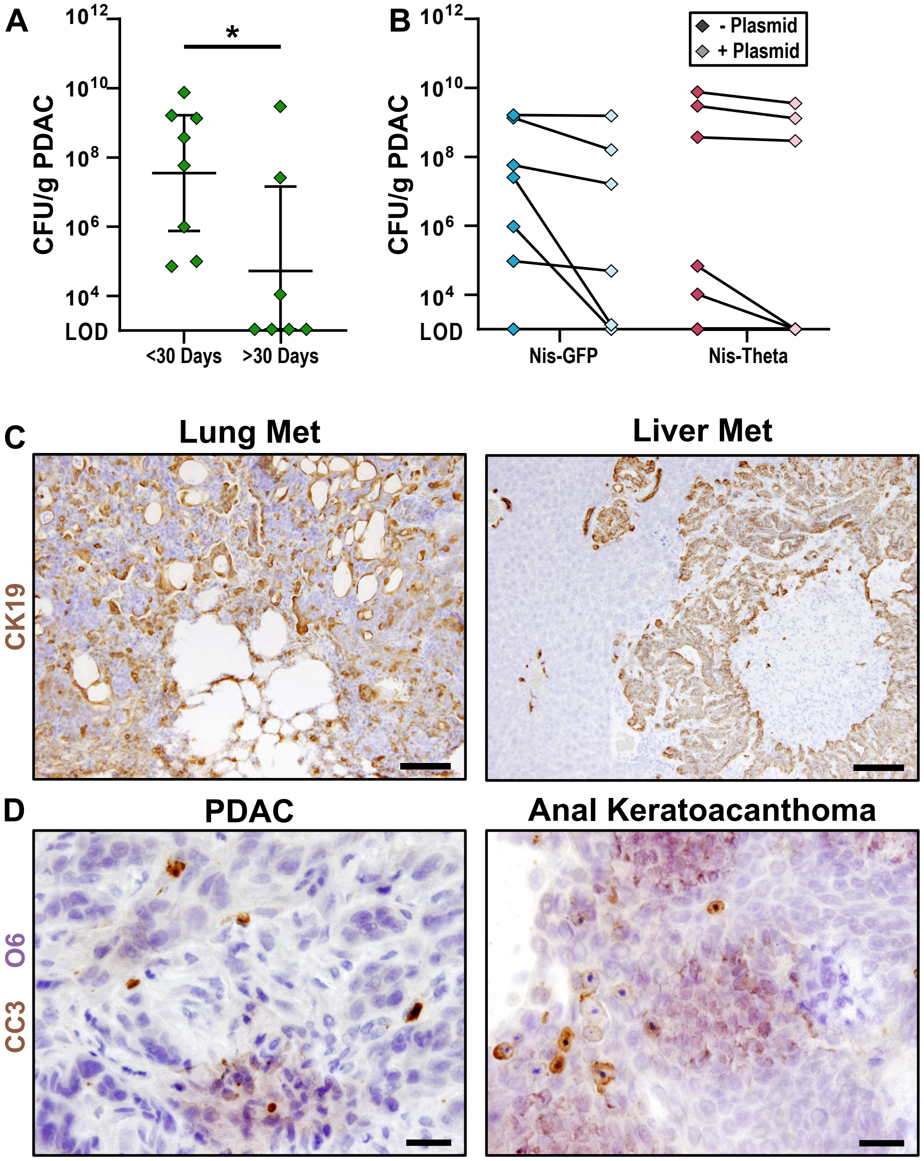


**Fig. S3. Tumor colonization dynamics of therapeutic bacteria.** (**A**) Endpoint biodistribution of Nislux bacteria in primary PDAC tumor separated by time on study. Data are presented as the geometric mean ± 95% CI. *$P$<0.05, Mann-Whitney, non-parametric t-test. (**B**) Percentage of Nislux bacteria recovered at endpoint that retained the AxeTxe stabilized therapeutic plasmid based on biodistribution on antibiotic selection plates. (**C**) Representative images of IHC for malignant epithelia (CK19, brown) identifying metastatic lesions in lung and liver tissues from bacteria treated mice. (20x magnification, scale bar = 100 µm). (**D**) Representative images of dual IHC for apoptosis (CC3, brown) and Nislux bacteria (O6, purple) demonstrating co-localization of bacteria with apoptotic cells. (100x magnification, scale bar = 20 µm).
