## Supplemental Figure 4 for "“Tumor-selective treatment of metastatic pancreatic cancer with an engineered, probiotic living drug”"

**Supp Fig. 4**

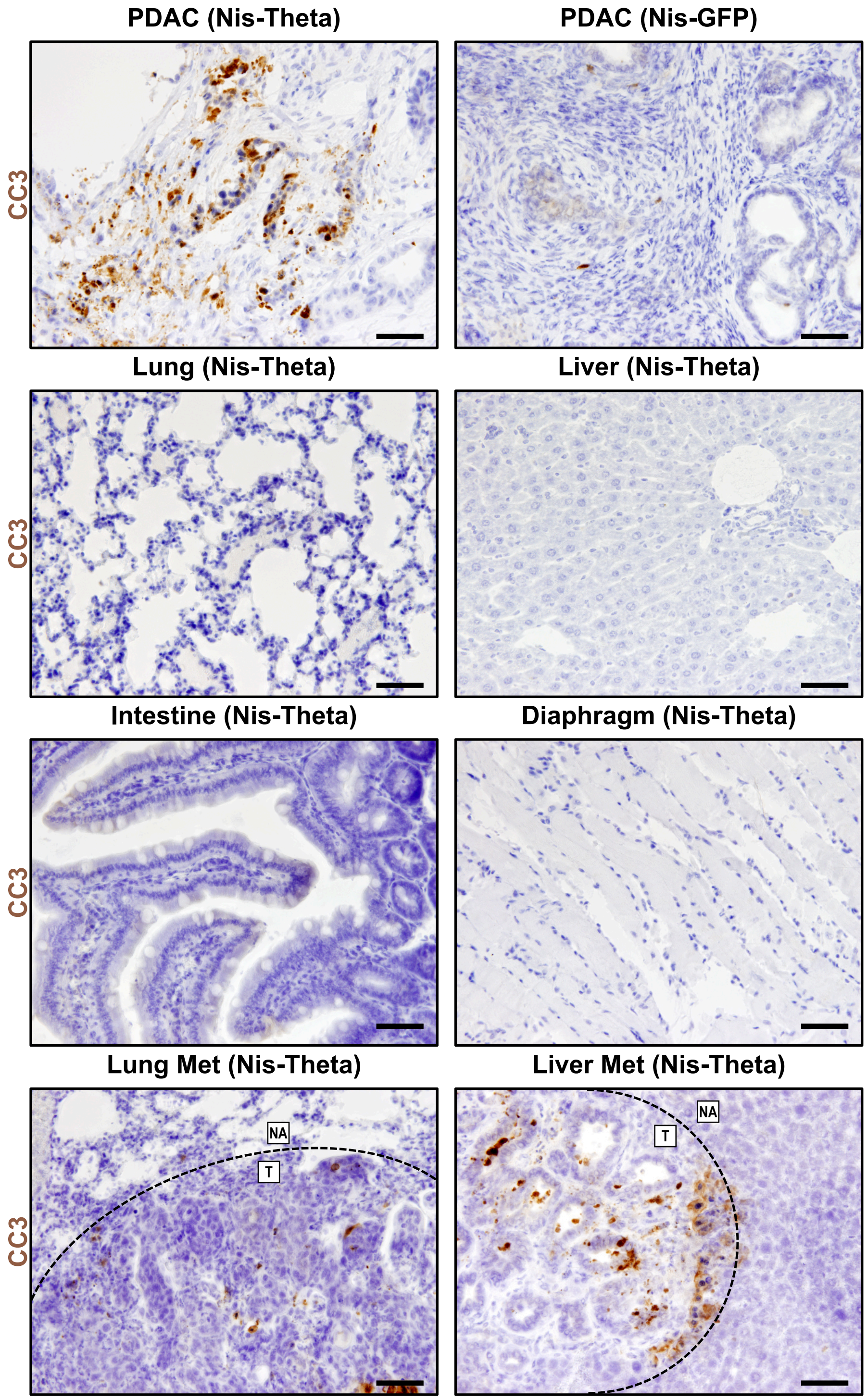


**Fig. S4. Evidence of cell death and tissue damage is limited to Nis-Theta treated tumor tissue, including metastatic lesions.** Representative images of IHC for apoptosis marker CC3 used for quantifying *in vivo* tissue damage in Fig. 3H. Dotted line indicates border between tumor (T) and normal adjacent (NA). (40x magnification, scale bar = 50 µm).
